# Functional Annotation Signatures of Disease Susceptibility Loci Improve SNP Association Analysis

**DOI:** 10.1101/000158

**Authors:** Edwin S. Iversen, Gary Lipton, Merlise A. Clyde, Alvaro N. A. Monteiro

## Abstract

We describe the development and application of a Bayesian statistical model for the prior probability of phenotype–genotype association that incorporates data from past association studies and publicly available functional annotation data regarding the susceptibility variants under study. The model takes the form of a binary regression of association status on a set of annotation variables whose coefficients were estimated through an analysis of associated SNPs housed in the GWAS Catalog (GC). The set of functional predictors we examined includes measures that have been demonstrated to correlate with the association status of SNPs in the GC and some whose utility in this regard is speculative: summaries of the UCSC Human Genome Browser ENCODE super–track data, dbSNP function class, sequence conservation summaries, proximity to genomic variants included in the Database of Genomic Variants (DGV) and known regulatory elements included in the Open Regulatory Annotation database (ORegAnno), PolyPhen–2 probabilities and RegulomeDB categories. Because we expected that only a fraction of the annotation variables would contribute to predicting association, we employed a penalized likelihood method to reduce the impact of non–informative predictors and evaluated the model’s ability to predict GC SNPs not used to construct the model. We show that the functional data alone are predictive of a SNP’s presence in the GC. Further, using data from a genome–wide study of ovarian cancer, we demonstrate that their use as prior data when testing for association is practical at the genome–wide scale and improves power to detect associations.

## 1 Introduction

The purpose of genetic association studies is to discover genetic loci that contribute to an inherited trait, identify the variants behind these associations and ascertain their functional role in determining the phenotype (Manolio, 2010). Modern association studies bring to bear on this problem high coverage genotype data, comprehensive databases of genetic variation that allow imputation of most common ungenotyped variants to high accuracy and extensive, publicly available, *in silico* resources housing a growing assortment of genomic data that allow functional characterization of vast regions of the human genome. In the typical genome–wide association study (GWAS), the first two forms of data are combined to reconstruct genotypes to a desired density and these genotypes are then systematically tested for association with the phenotype. The functional annotation data are most frequently used in *post hoc* interpretation of evident associations raised by the analysis (Freedman et al., 2011).

To date, functional annotation data have rarely played more than an indirect role in assessing evidence for association. For example, they may be used to suggest candidate genes and SNPs for study or to support links between candidate SNPs and genes. While methods to incorporate functional annotation data *a priori* in genetic association analyses exist, they are infrequently used. The prevailing approach to this is via a two–staged hierarchical model in which coefficients in the stage I generalized linear model for phenotype given genotype and exposure measurements are regressed, in stage II, on the annotation data (Witte et al., 1994; Aragaki et al., 1997; Hung et al., 2004, 2007). This is limited to analysis of a modest number of variants and does not make use of prior data derived from previous association studies to inform the nature of that relationship.

It is becoming increasingly clear that a widening array of annotation data correlates with a variant’s having been associated with a human phenotype (Hindorff et al., 2009; Nicolae et al., 2010; The ENCODE Project Consortium, 2012; Schaub et al., 2012). In what follows, we describe a formal approach to inference for association that combines functional annotation data (through a prior distribution) with genotype data (through a sampling model for the phenotype given genetic and other covariate data). We construct the prior distribution through careful analysis of SNPs housed in the GWAS Catalog (Hindorff et al., 2009). We refer to the linear combination of the annotation variables defined by this model and evaluated for a given SNP as its ‘functional annotation signature.’ We show that functional signatures so derived are predictive of the association status of SNPs not used in their creation and that, when coupled with genetic association data following the method we describe, improve the efficiency of association testing in a GWAS study of ovarian cancer.

## 2 Results

The ultimate goal of association studies is to identify the set of common polymorphisms that influence a phenotype. This goal is approached through a statistical analysis designed to measure the evidence in favor of association followed by a decision rule used to declare each variant’s true status as ‘associated,’ ‘uncertain,’ or ‘unassociated.’ The data that inform these analyses usually comprise phenotype labels, SNP genotype data and a set of non–genetic covariates in addition to functional annotations of the variants under study. The statistical analysis may take many forms, varying according to choice of modeling approach and inferential paradigm (Frequentist or Bayesian). The approach we develop here relies on Bayesian inference but can also be applied when the genetic association summaries are p-values. In this paradigm, prior data on a quantity of interest (such as the binary association status of a genetic variant) are updated to reflect evidence in the current data set.

A Bayesian analysis of genetic association data returns an estimate of the odds of association of each marker given the available data. When the data take two distinct forms — here subject–level phenotype, genotype and covariate data and variant–level functional annotations — the odds of association may be calculated in two stages, either by incorporating functional data prior to or following evaluation of the genetic data. The latter represents the heuristic typically followed in practice, whereby functional data is evaluated in an informal way (from the probabilistic point of view) conditional on evidence for association. Here we describe a model–based framework for combining functional and association data following the second factorization. We focus on the case–control study design for purposes of illustrating integration of the *a priori* (to association data) models for functional annotation data we describe below into analyses of genetic association data. Details of the models and their assumptions are provided in Methods.

When the functional data are incorporated as prior information, the odds of a SNP’s association given the functional and subject–level data can be written as the product of the Bayes factor (BF) in favor of association and the prior odds of association given the functional data. The BF is the ratio of the integrated likelihood of the phenotype data given the covariate and genotype data assuming the SNP is associated to the integrated likelihood of the phenotype data given the covariate data only (i.e. assuming the SNP is not associated). It is a commonly used Bayesian statistical measure of association and is calculated by the SNPTEST (Marchini et al., 2007) and BIMBAM (Servin and Stephens, 2007) packages for analysis of GWAS data. Alternately, Sellke et al. (2001) show that an upper bound on the Bayes factor in favor of association is approximately equal to −1/(*ep* log_*e*_(*p*)) when *p* < (1/*e*) and 1.0 otherwise, where *p* is the p-value for association. This allows the method to be used in conjunction with standard frequentist association testing software.

In short, the functional annotation data are incorporated into an analysis by formally updating the prior odds of association given the annotation data by a standard measure of genetic association. This process is depicted schematically in Figure 1. In what follows, we describe the model used to calculate prior odds of association and demonstrate its use in a GWAS of ovarian cancer. In it, the log of a SNP’s prior odds of association, its ‘functional signature,’ is a linear combination of the functional data.

**Figure 1.**
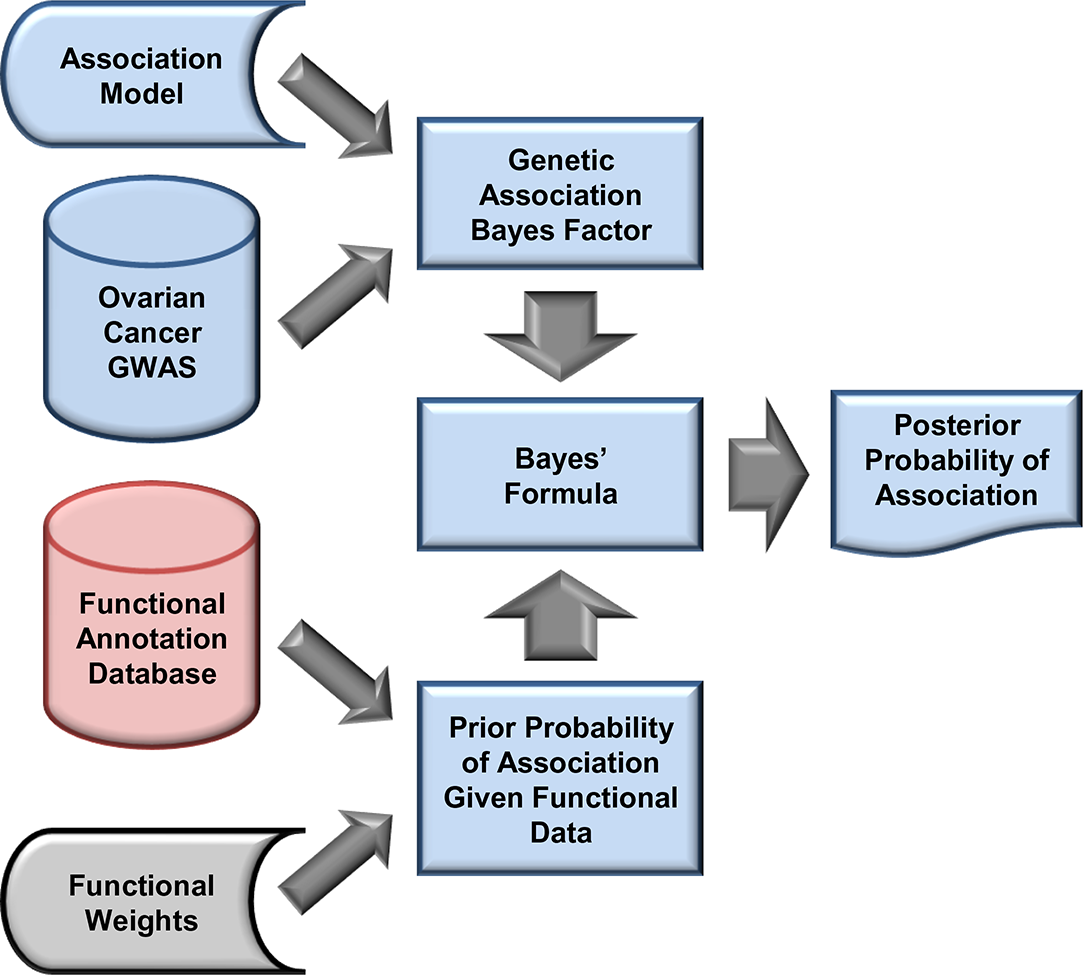
Two–staged procedure for integrating variant–level functional annotation data with subject–level genetic association data. At the first stage, functional annotation data are combined to estimate the prior (to observing the genetic association data) probability of association for each variant. At stage two, these estimates are combined with the Bayes Factor (a metric of association) in favor of genetic association via Bayes’ formula to estimate the posterior (to observing the functional and genetic association data) probability of association for each variant.

### 2.1 Functional Signatures of Known Associations

We constructed the functional annotation signatures by estimating the multivariate relationship between a set of functional annotation variables and the binary association status of a set of SNPs. Figure 2 provides a schematic of our approach. In brief, we identified a set of associated SNPs and, for each, we chose a matching, unassociated ‘control’ SNP. We divided the matched pairs into ‘training’ and ‘validation’ sets and used the former to construct a series of models to predict association status given the function data and used the latter to compare the performance of these models. We chose the model that demonstrated the best predictive accuracy in the validation data, as measured by concordance index, to define the functional annotation signatures.

**Figure 2.**
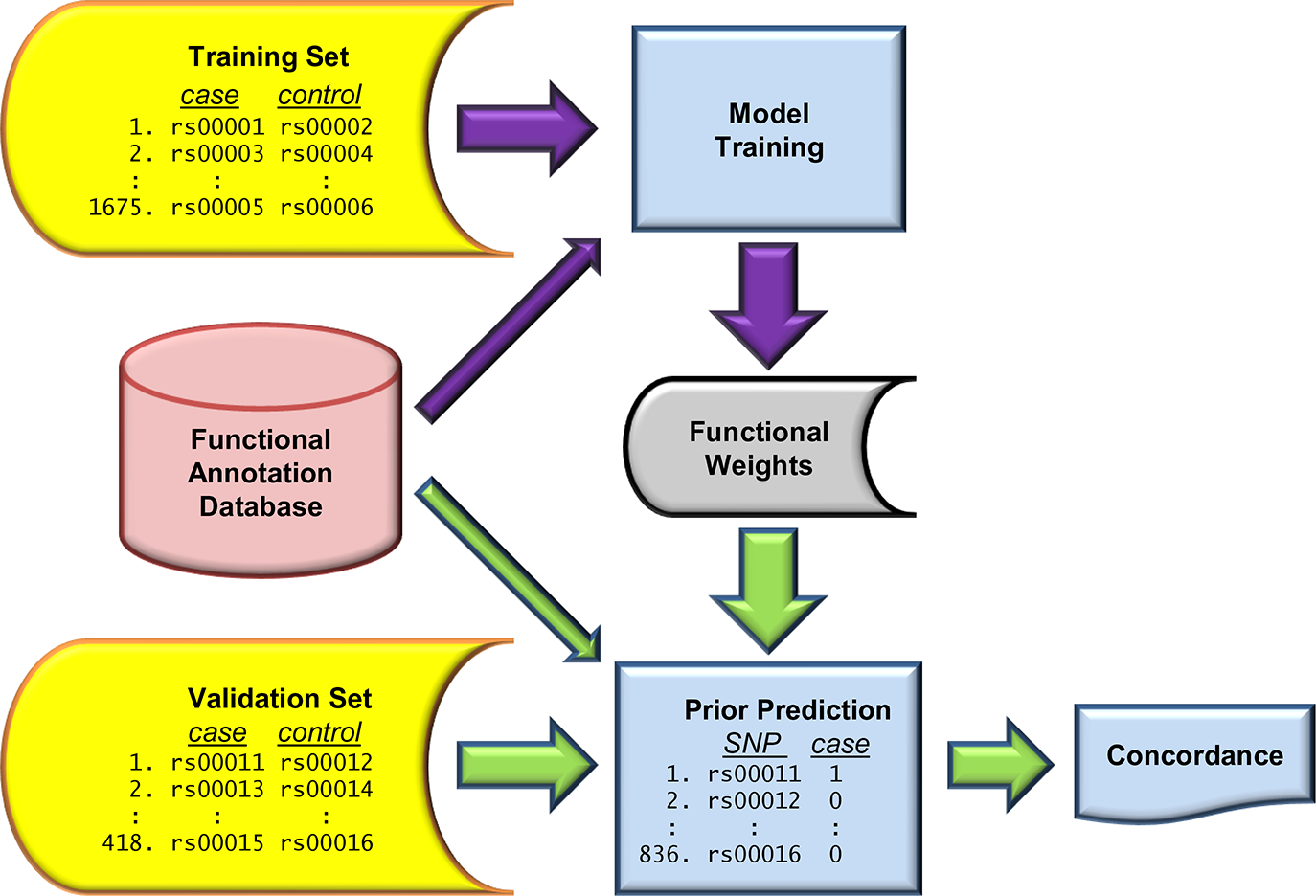
Construction and evaluation of models for (prior) probability of association given the functional annotation data. The purple arrows represent model construction (‘training’), while the green arrows represent evaluation of the models. Construction of the training set, validation set and functional annotation database are depicted in Figure 4 and described in Methods. The training data were used to construct a series of models, each distinguished by the coefficients (or ‘weights’) it assigns to the various annotation variables. We chose the best amongst these by comparing their predictions in the validation set using the concordance index.

We began by constructing a matched case–control study *of SNPs* in which the cases were drawn from the GWAS Catalog (Hindorff et al., 2009) and the controls were identified from the HapMap database, Release 27, Phases II and III merged genotypes. We identified 2,093 case SNPs and, for each, identified one control SNP matched on chromosome, minor allele frequency and the genotyping platform(s) it appeared on. Since SNPs in the GWAS Catalog are arguably more frequently tags than the directly associated variant, we followed Hindorff et al. (2009) and identified ‘LD partners’ for each case and control SNP. We grouped each case and control SNP together with its LD partners to form blocks.

Using on–line bioinformatics resources, we assembled a set of functional annotation variables representing a variety of contextual descriptions and empirical measurements with which we annotated each of the 48,889 case, control and LD partner SNPs. We included annotation variables shown to be correlated with presence in the GWAS Catalog or that we believed likely to be so. These were: dbSNP function designation; summaries of ENCODE Project (The ENCODE Project Consortium, 2007, 2011) data on transcription levels assayed by RNA–seq (Mortazavi et al., 2008; Langmead et al., 2009), measures of signal enrichment for H3K4Me1, H3K27Ac and H3K4Me3 histone modifications associated with enhancer and promoter activity (Bernstein et al., 2006; Mikkelsen et al., 2007), evidence for overlap with a DNaseI hypersensitivity cluster (Sabo et al., 2006, 2004) and evidence for transcription factor binding (Euskirchen et al., 2004, 2007; Martone et al., 2003; Robertson et al., 2007; Rozowsky et al., 2009); PhyloP evolutionary conservation scores (Siepel et al., 2006); indicators for whether or not the variant falls in a region of known copy number variation, a region containing insertions or deletions or a region with inversions (Iafrate et al., 2004; Zhang et al., 2006); PolyPhen–2 (Adzhubei et al., 2010) probability that a mutation is damaging; and RegulomeDB score (Boyle et al., 2012). The latter represents a synthesis of regulatory data derived from ENCODE and other sources. While not a comprehensive set, they covered the major annotation classes available at the time of analysis and are readily available to individuals executing an association study. The infrastructure and methods described here are easily updated to accommodate new variables as they become generally available. Table 1 lists the 57 variables that we used to construct the functional signatures of association.

**Table 1.**
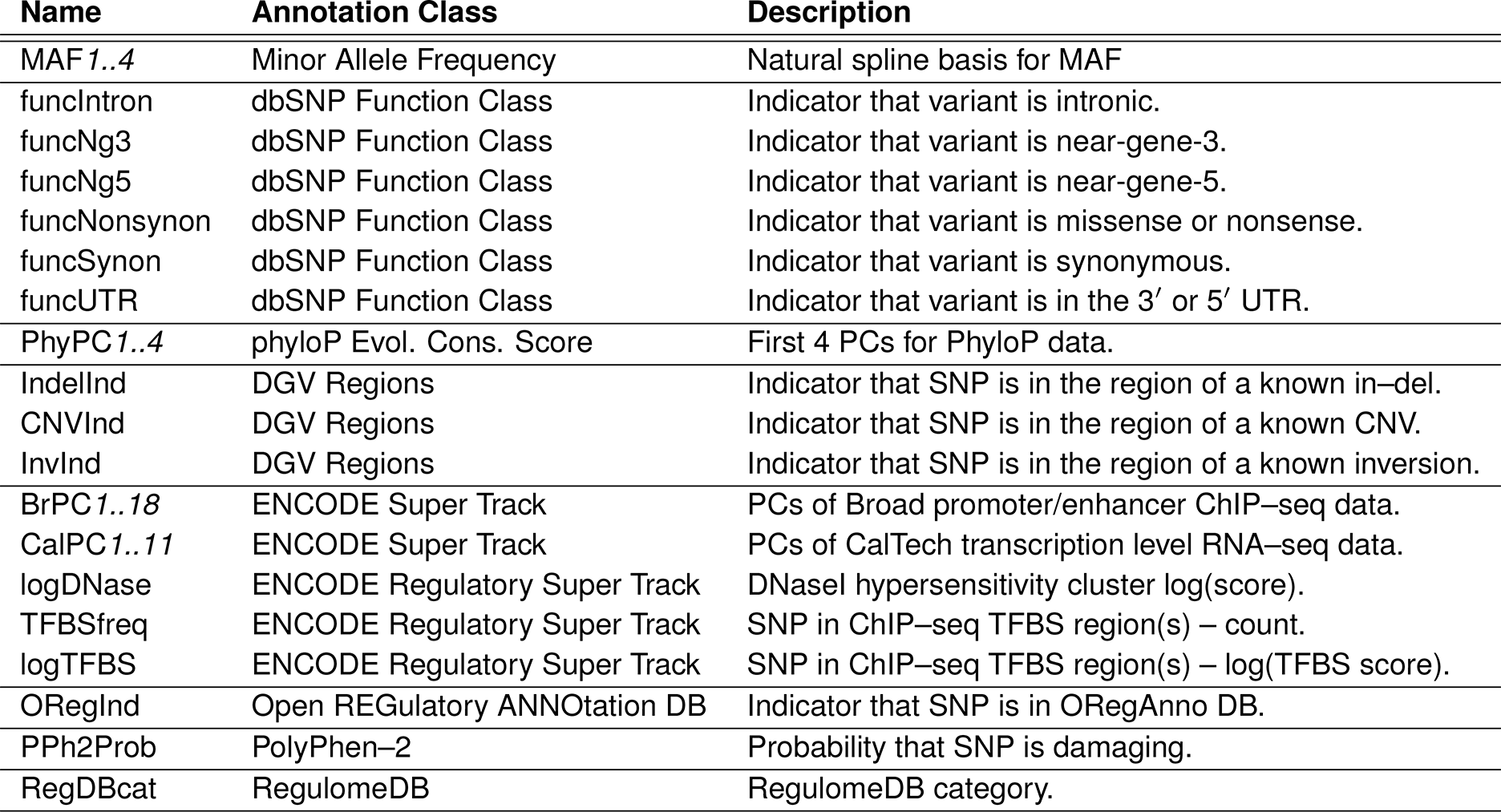
Annotations used to construct the functional signatures. Definitions of the 54 variables appearing in the prior model for association status arranged by type/class of annotation.

The 48,889 SNPs included in the analysis were grouped into 2,093 case and an equal number of control blocks. We randomly selected 1,675 of these matched case–control pairs for development of the model (the ‘training set’) and left the remaining 418 pairs for model evaluation and comparison (the ‘evaluation set’). We modeled the probability that a SNP is associated given its functional data using a logistic regression model. Further, we assumed that each case block contained one or more associated SNPs and that each control block contained none.

While the assembled list of functional predictors includes measures that have been demonstrated to correlate with the association status of SNPs in the GC, it also includes a number of measures whose utility in this regard was unclear. Hence, we expected that only a fraction of the 57 variables would contribute to predicting phenotype association. We used shrinkage priors (Hans, 2009; Richardson et al., 2011) to reflect this belief and chose the normal–exponential–gamma (NEG) distribution for its ability to penalize heavily weakly determined predictors and to penalize weakly those that are well determined (Griffin and Brown, 2007; Hoggart et al., 2008; Griffin and Brown, 2010). Further details of the model and the Markov chain Monte Carlo (MCMC) algorithm used for inference can be found in Methods.

Table 2 provides a summary of the coefficient estimates obtained for the binary regression of association status on the 57 functional annotation variables. Because all variables in the model were standardized, coefficients measure the difference in the log–odds of phenotype association attributed to an increase of one standard deviation in the covariate when the others remain fixed. The majority of predictive variation (51%) in the functional scores as measured in the control block SNPs from the validation set, is due to the Broad promoter/enhancer ChIP–seq principal components (PCs) and nearly all (*>* 97%) of this variation is due to PCs 1, 2, 4, 5, 6, 8 and 13. Each PC is a linear combination of the 75 summary statistics of the 25 assays. Supplemental Figure 1 depicts the loadings (weights in the linear combinations) for these PCs as they depend on histone modification, cell line and summary statistic. Grossly, PC 1 measures total signal strength across all cell lines and histone modifications, PC 2 contrasts average signal strength of the H3k4me3 assay with variation over all assays, while the remaining PCs each contrast signal in one subset of cell lines with that in another (PC 4: HMEC and NHEK *versus* GM12878 and K562; PC 5: GM12878, HMEC and NHEK *versus* HSMM, HUVEC and NHLF; PC 6: H1-hESC, HepG2 and HSMM *versus* GM12878 and HUVEC; PC 8: K562 *versus* H1-hESC; and PC 13: HepG2 *versus* H1-hESC).

**Table 2.** Summary of estimates for the model for association status given the functional annotation data. Estimates of the posterior mean and standard deviation are provided for each coefficient in the model along with the ratio of these quantities, a ‘signal-to-noise’ measure analogous to the Z statistic.

| **Coefficient** | **Mean** | **SD** | **Mean/SD** | **Coefficient** | **Mean** | **SD** | **Mean/SD** |
| --- | --- | --- | --- | --- | --- | --- | --- |
| MAF1 | 0.029 | 0.0272 | 1.051 | CalPC8 | 0.096 | 0.0608 | 1.584 |
| MAF2 | 0.003 | 0.0185 | 0.151 | CalPC9 | 0.009 | 0.0218 | 0.406 |
| MAF3 | 0.018 | 0.0236 | 0.759 | CalPC10 | -0.019 | 0.0234 | -0.809 |
| MAF4 | -0.008 | 0.0199 | -0.425 | CalPC11 | -0.044 | 0.0468 | -0.943 |
| BrPC1 | -0.348 | 0.0388 | -8.983 | PhyPC1 | 0.225 | 0.0452 | 4.982 |
| BrPC2 | 0.174 | 0.0360 | 4.845 | PhyPC2 | -0.023 | 0.0308 | -0.742 |
| BrPC3 | 0.002 | 0.0172 | 0.099 | PhyPC3 | 0.053 | 0.0395 | 1.336 |
| BrPC4 | -0.077 | 0.0301 | -2.561 | PhyPC4 | 0.024 | 0.0289 | 0.839 |
| BrPC5 | 0.149 | 0.0301 | 4.932 | funcIntron | -0.003 | 0.0199 | -0.160 |
| BrPC6 | 0.097 | 0.0329 | 2.961 | funcNg3 | 0.002 | 0.0238 | 0.098 |
| BrPC7 | -0.019 | 0.0229 | -0.825 | funcNg5 | 0.003 | 0.0200 | 0.158 |
| BrPC8 | -0.078 | 0.0312 | -2.498 | funcNonsynon | 0.089 | 0.0387 | 2.308 |
| BrPC9 | -0.012 | 0.0202 | -0.573 | funcSynon | -0.007 | 0.0236 | -0.283 |
| BrPC10 | 0.007 | 0.0182 | 0.407 | funcUTR | 0.002 | 0.0210 | 0.078 |
| BrPC11 | 0.021 | 0.0239 | 0.887 | logDNase | 0.011 | 0.0253 | 0.419 |
| BrPC12 | -0.039 | 0.0290 | -1.343 | TFBSfreq | 0.024 | 0.0267 | 0.885 |
| BrPC13 | -0.094 | 0.0318 | -2.972 | logTFBS | 0.019 | 0.0299 | 0.641 |
| BrPC14 | -0.000 | 0.0175 | -0.015 | ORegInd | 0.027 | 0.0236 | 1.163 |
| BrPC15 | -0.039 | 0.0286 | -1.354 | IndelInd | -0.023 | 0.0337 | -0.693 |
| BrPC16 | 0.009 | 0.0185 | 0.467 | CNVInd | 0.059 | 0.0321 | 1.842 |
| BrPC17 | -0.015 | 0.0213 | -0.696 | InvInd | 0.090 | 0.0305 | 2.938 |
| BrPC18 | -0.014 | 0.0210 | -0.688 | rDBcat1 | 0.014 | 0.0257 | 0.550 |
| CalPC1 | -0.103 | 0.0492 | -2.084 | rDBcat2 | 0.066 | 0.0378 | 1.741 |
| CalPC2 | 0.090 | 0.0441 | 2.030 | rDBcat3 | 0.012 | 0.0264 | 0.467 |
| CalPC3 | -0.019 | 0.0270 | -0.709 | rDBcat4 | 0.116 | 0.0461 | 2.508 |
| CalPC4 | 0.086 | 0.0457 | 1.889 | rDBcat5 | 0.106 | 0.0527 | 2.003 |
| CalPC5 | 0.053 | 0.0395 | 1.350 | rDBcat6 | -0.056 | 0.0552 | -1.018 |
| CalPC6 | -0.012 | 0.0233 | -0.496 | pph2prob | 0.078 | 0.0269 | 2.905 |
| CalPC7 | 0.002 | 0.0207 | 0.078 |  |  |  |  |

The sequence conservation PCs collectively make the next largest contribution, explaining 16% of variation in the functional scores; PCs 1 and 3 explain *>* 97% of this. Each PC is a linear combination of the summary statistics of the 28 and 44 species PhyloP scores, each for all species and restricted to placental mammals. Supplemental Figure 2 graphs the loadings for these PCs as they depend on number of species, depth of alignment and summary statistic. Briefly, PC 1 measures total signal strength across scores with the scores based on the 28-way alignment weighted more heavily than those based on the 44-way alignment, while PC 3 contrasts the 28-way with 44-way scores.

The CalTech RNA–seq PCs collectively explain 10% of the signature, with PCs 1, 2, 4 and 8 contributing 87% of this. Supplemental Figure 3 depicts the loadings for these PCs as they depend on cell line and summary statistic. PC 1 provides a measure of total signal strength across all cell lines, while the remaining PCs each contrast signal in one subset of cell lines with that in another (PC 2: H1-hESC and K562 *versus* GM12878 and NHEK; PC 4: GM12878 and H1-hESC *versus* K562, NHEK and HepG2; PC8: HUVEC *versus* NHEK).

RegulomeDB score explains the next largest fraction (8%) of variation. It is represented by six variables, each indicating a functional category; category 7 serves as the reference (’baseline’). Categories 2, 4, 5 and 6 explain 99% of this variation suggesting that other annotation variables in the model better characterize the probability of phenotype association for variants in categories 1 and 3. Virtually all (96%) of the 2.5% contribution to variation made by the DGV variables is due to the copy number and inversion variables. Finally, the dbSNP functional class variables are the only remaining that contribute more (=1.7%) than 1% of the variation in functional scores. Virtually all (99%) of this contribution is due to the non–synonymous designation within which the PolyPhen–2 probability contributes significant resolution to the model.

We estimated the concordance indexes (equivalent to AUC, area under the ROC curve) for each model using the 418 matched case–control block pairs in the validation set as a tool for comparing the accuracy of their out–of–sample predictions. Table 3 provides the estimates of concordance and associated 95% interval estimates. While the concordance statistics are not discernibly different from one another, the best out–of–sample predictive ability is achieved using the model with the prior distribution having the strongest shrinkage properties, i.e. the ‘NEG3’ model.

**Table 3.**
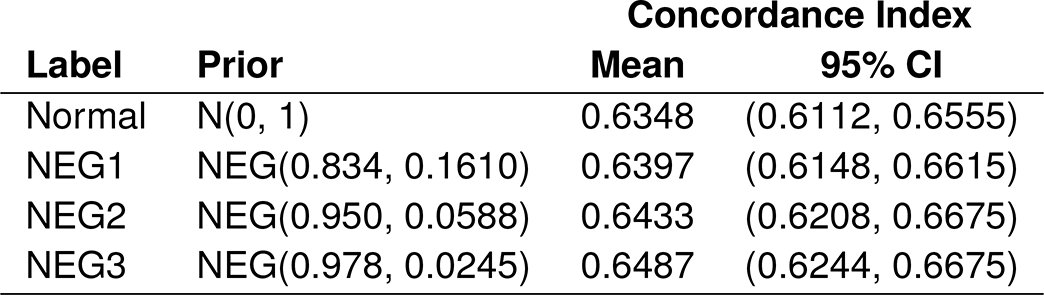
Means and 95% interval estimates of the concordance indices for each of the four models.

### 2.2 Application to an Ovarian Cancer Multi–GWAS Study

Here we compare the ranks assigned to a group of variants in a GWAS analysis when those ranks are calculated with and without the functional annotation data. Each in the group of variants is assumed to have known association status (associated/unassociated) with epithelial ovarian cancer, where this determination is based on confirmatory studies subsequent to the GWAS. The group is constructed as follows. There are currently 11 published, genome–wide significant loci for epithelial ovarian cancer. Nine of the 11 have come to light through analysis of genome–wide SNP data. These are rs3814113 (Song et al., 2009), rs8170 (Bolton et al., 2010), rs2072590, rs2665390, rs7814937, rs9303542 (Goode et al., 2010), rs11782652, rs7084454, rs757210 (Pharoah et al., 2013). The remaining two (rs10069690 and rs2077606) were identified by candidate gene/pathway investigations (Bojesen et al., 2013; Permuth-Wey et al., 2013); all 11 have been evaluated in very large confirmatory studies. We consider these to be ‘true positive’ variants. Our analysis of data from the large–scale follow–up study of GWAS candidates described in Pharoah et al. (2013) allowed us to identify a group of variants with strong evidence *against* association that we treat here as ‘true negatives.’

Table 4 summarizes the GWAS results for the true positive and true negative SNPs when the analysis is conducted with (subscript ‘A + F’) and without (subscript ‘A’) the functional signatures and where the association summaries (Bayes factors) are calculated directly (columns labeled ‘Bayesian Analysis’) and approximated from the results of standard likelihood ratio tests using the method of Sellke et al. (2001) (columns labeled ‘P-Value Approximation’). We focus here on results of the Bayesian analysis, noting that the approximate method yields very similar results. Note that the candidate SNPs are ranked substantially lower than the GWAS ‘hits.’ Indeed, the evidence in the association data related to these variants is actually *against* association (both of their Bayes factors are less than 1.0). The GWAS hits are all ranked in the top 50,000 (of approximately 2.5 million) by the same measure and all have Bayes factors of at least 3 to 1 in favor of association.

**Table 4.**
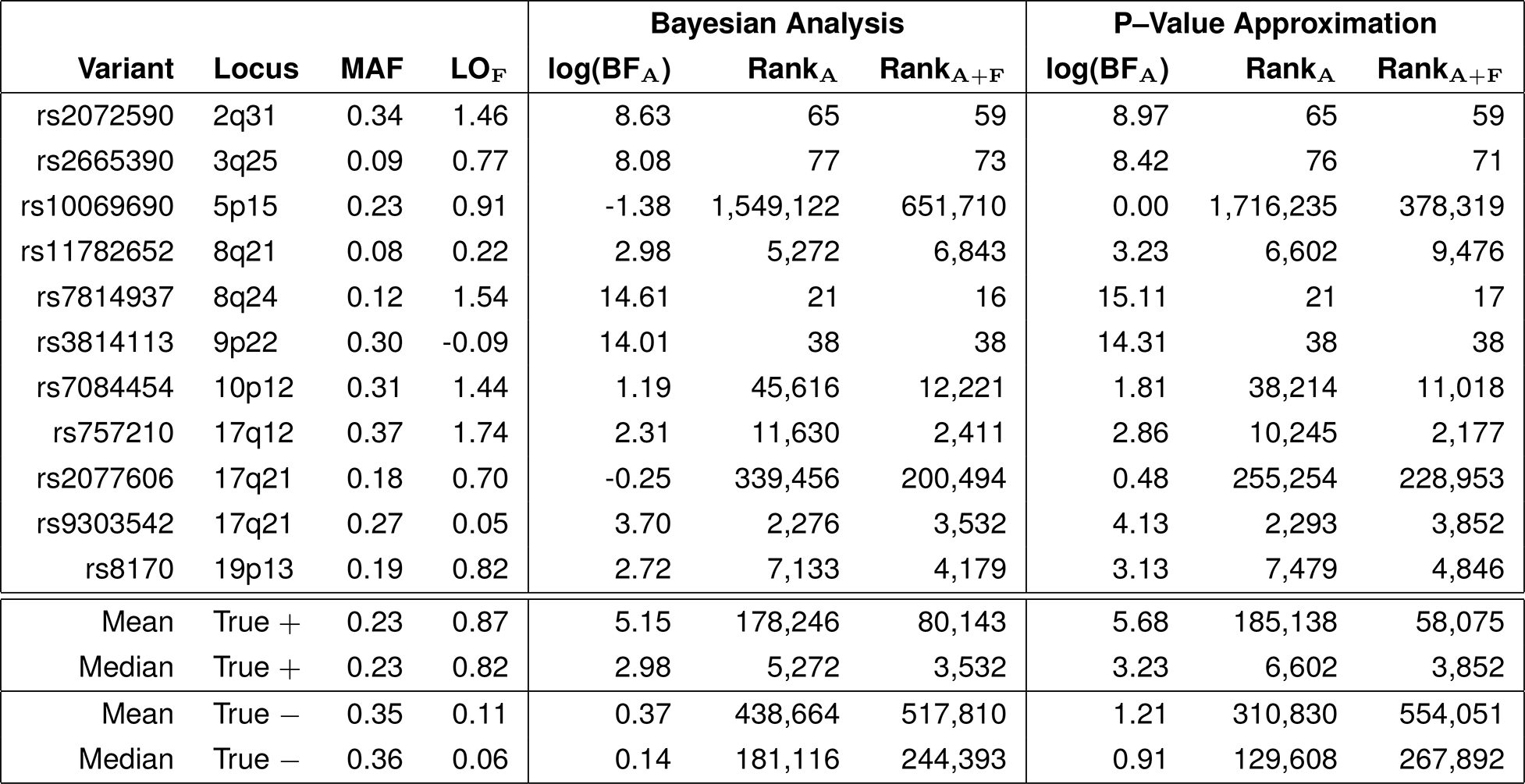
Functional signatures improve inference for association status in a GWAS of ovarian cancer. Ranks of known associated variants (labeled ‘true +’) tend to improve (i.e. are closer to one) when association and functional data are incorporated in the analysis (Rank_A+F_) relative to when only the association data are used (Rank_A_) and, hence, are more likely to be studied further. Conversely, ranks of (very likely) unassociated variants (labeled ‘true −’) tend to fall with inclusion of the functional data. The functional data for a given variant is summarized by its ‘functional signature,’ defined as the prior log–odds of its association given the functional data (LO_F_). Aggregate (mean and median) values are provided for the true + set and the true − set. Results are provided both for when the Bayes Factors in favor of genetic association (BF_A_) are estimated from a Bayesian analysis and for when they are approximated using a transformation of p-values. Ranks are out of approximately 2.5M variants.

Only two of the truly associated SNPs (rs11782652 and rs9303542) are ranked higher when the functional data are ignored than when they are used, however their respective changes in rank are small. The median (alt. average) rank of the truly associated SNPs was 5,272 (178,246) without and 3,532 (80,143) with the functional data included. If design constraints allowed only for followup of the top 5,000 variants, a larger fraction (7/11) would be discovered with addition of the functional data than without (5/11); with followup of 10,000 variants, these fractions become 8/11 and 7/11. In contrast, when the function data were included the median (alt. average) rank among a set of ‘true negative’ SNPs increased from 181,116 (438,664) to 244,393 (517,810), while the number selected for followup fell from 244 to 204 under the 5K scenario and from 443 to 373 under the 10K scenario.

#### Functional signatures of tag SNPs correlate with function of tagged SNPs

While a few of the functional variables, such as the function class designation ‘nonsynonymous,’ incorporated in the signature are base pair specific, most map to contiguous regions of 100’s or 1000’s of base pairs. Hence, the functional signatures associated with nearby SNPs are correlated. Figure 3 is a plot of the correlation between the functional signatures of adjacent SNPs that passed QC in the ovarian cancer GWAS described above as a function of the distance, measured in base pairs (BPs), between the two variants. This correlation is greater than 0.72 (alt 0.68) for more than 80% (alt 97.5%) of adjacent variants, corresponding to those at distances of 1470 (alt 4376) BPs or less. Hence, while there are gains to be realized in doing so, it is not necessary to impute to and annotate at the highest possible density to realize an increase in power to detect association through the use of functional signatures, a fact we demonstrated empirically above. Note that typical BP distances between tagged (not genotyped or imputed) variants and their nearest tag will be on the order of one half of the distances reported here for adjacent tags.

**Figure 3.**
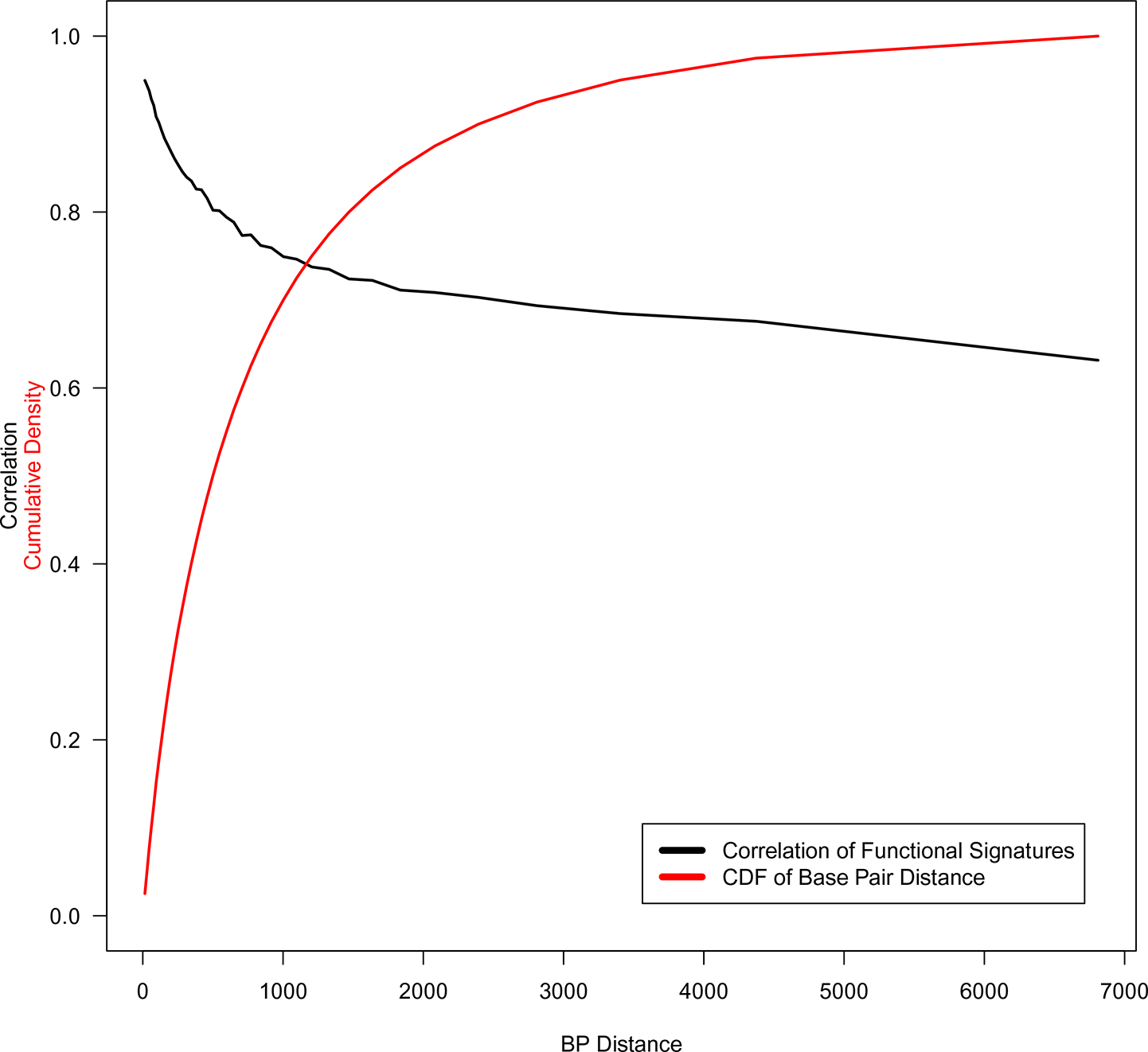
Correlation of functional signatures between adjacent HapMap II/III SNPs as a function of base pair distance (black line). Cumulative distribution function(CDF) of base pair distances across the genome (red line).

## 3 Discussion

Using the GWAS Catalog as a sampling frame, we developed a model for the probability that a given polymorphism is associated with an observable human phenotype given a set of functional annotation variables and demonstrated that this model has the ability to predict a set of phenotype associated variants not used in the model building exercise. We demonstrate several methods for incorporating functional annotation signatures defined by this model and evaluated for a SNP’s annotation data as prior data and show through example that by doing so we improve the efficiency of GWAS scale analysis to identify true positive associations for follow-up study.

The approach we describe is computationally tractable and scalable to modern genome–wide analysis. Our use of penalized regression techniques to model the functional data and construct the function signatures allows us to consider a relatively large number of individual annotation variables while controlling for over–fitting. We evaluated sensitivity of the model’s out-of-sample predictions to choice of shrinkage prior and found that the most aggressive choice we examined, the model whose results are summarized herein, resulted in the best out-of-sample concordance estimates. Our approach can be expanded and adapted to incorporate more detailed annotation data such as was recently released by the ENCODE consortium (The ENCODE Project Consortium, 2012) or generated experimentally in individual labs.

In principle, estimates of the parameters in the model for SNP association status given the functional data can be refined via Bayesian updating as part of an association analysis. This requires an additional layer of analysis that is feasible, but computationally demanding to implement on a genome–wide scale. However, the value of this will be limited in settings where there are few truly associated SNPs and/or the case–control data supporting associations are weak, i.e. the vast majority of applications. Here, Bayesian updating will yield estimates equivalent to those using the approach we describe above up to Monte Carlo simulation error. Indeed, we formally compared the two approaches using the ovarian cancer GWAS data and found little change in the median ranks of the true positive (3,532 *versus* 3,705) and true negative SNPs (244,393 *versus* 248,459). This suggests that the added value of Bayesian updating to the functional signatures will typically be limited.

Performance for our integrative approach likely depends on the depth, specificity and density of coverage of the available annotation data. The current study defines a starting point and benchmark in each of these dimensions. In particular, while the depth of annotation considered here is sufficient to noticeably improve inference for association, it is clear from recent ENCODE Project Consortium publications that it reflects only a small fraction of the complexity present in the regulatory landscape. Further, none of the annotation variables are tailored to the outcome phenotype; indeed, the ENCODE super track data enter the model through linear combinations of the cell–line specific measurements, effectively averaging over cell type. Many regulatory processes are cell–type–specific (The ENCODE Project Consortium, 2012; Schaub et al., 2012) and hence will be more informative for a given phenotype if measured in the appropriate cell type. However, determining the relevant annotation data, assuming it exists, for a given phenotype requires domain expertise and more careful modeling to create functional signatures. While Bayesian updating did not improve inferences in the ovarian cancer GWAS example, a generalization of it that couples the existing signature structure with context–specific annotations such as cell type specific eQTL data and an independent prior distribution on its multivariate adjusted effect is one approach to improving specificity.

Finally, our analyses have been carried out entirely at the HapMap III density. Our approach succeeds at this density because the functional signatures of SNPs nearby, at distances typical of HapMap III, are highly correlated and hence the functional signatures of HapMap III polymorphisms essentially tag function of nearby polymorphisms not in the database. As coverage (genotype/imputation density) of the typical association study becomes more complete, the need to rely on correlations between functional signatures will diminish and their power to assist in identifying and localizing associations is expected to increase. Association analyses at the density of the 1000 Genomes Project database (The 1000 Genomes Project Consortium, 2012) are now possible and will likely become common. The specificity of the functional signatures should improve when reconstructed and applied at this density as we plan to do as we continue to develop this approach.

## 4 Methods

### 4.1 Association Analysis Given Annotation Data

Let **G** be an *n* by *p* matrix of SNP genotypes, *D* be an *n* by 1 vector of disease indicators where *D*_*i*_ = 1 if individual *i* has the disease and *D*_*i*_ = 0 otherwise, **X** be an *n* by *r* matrix of covariates used in the association model and **F** be a *p* by *m* matrix of SNP–level functional annotation data where *n* is the number of individuals, *p* is the number of SNPs, *r* is the number of covariates and *m* is the number of annotation variables. Finally, let *A* be a *p* by 1 vector of 0–1 indicators of the (unknown) association status of the variants where *A*_*s*_ = 1 if SNP *s* is associated with the phenotype of interest.

In what follows, we specify the likelihood for the association indicator given the association (**X**, *D*, *G*) and function (**F**) data. To this end, we let Pr(*A* | *D*, **X**, **G**, **F**) 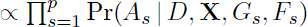. This relies on two assumptions: (1) that the *A*_*s*_’s are conditionally independent given (**X**, *D*, **G**, **F**) and (2) that the *A*_*s*_’s are conditionally independent of other variants (**G**_−*s*_, **F**_−*s*_) given (**X**, *D*, *G*_*s*_, *F*_*s*_). The notation **G**_−*s*_ indicates the matrix obtained by removing column *s* from **G**.

Further, we assume that the disease phenotype data are conditionally independent of the functional data for SNP *s* given the association status of that SNP, the covariate data and the genotype data for that SNP and that the association status indicator for SNP *s* is conditionally independent of the covariate data and its genotype data given its functional data. The latter assumption may be violated, for example, if the genotype data *G*_*s*_ carries information about function (e.g. minor allele frequency) not included in **F**. Given this, the odds of association of SNP *s* given its association and functional data can be written as the product of the (prior) odds of its association given its functional data times the (integrated) likelihood ratio or Bayes Factor (BF) of the phenotype given the SNP genotype and other covariate data, i.e.

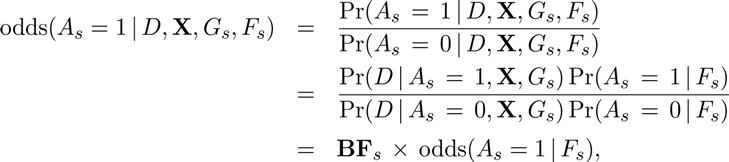

We describe estimation of the association summary Bayes factor below.

Given the binary, logistic link model developed below for association status given the functional data and the parameters *α* and *β*, odds(*A*_*s*_ | *F*_*s*_) = exp (*α* + *F*_*s*_*β*) and hence, given *α* and *β*

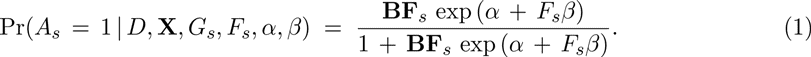

Provided that estimates of *α* and *β* are available from an external analysis such as described in the next section, one can estimate Pr(*A*_*s*_ = 1 | *D*, **X**, *G*_*s*_, *F*_*s*_) by

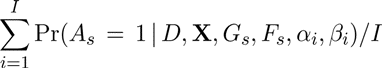

where the *α*_*i*_ and *β*_*i*_ are samples from the posterior distribution from an analysis such as described in Section 2.1.

The above procedure depends on estimates of the marginal likelihoods,

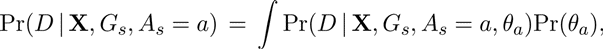

of the association data for each SNP under H_o_(*A*_*s*_ = 0) and under H_a_(*A*_*s*_ = 1). Pr(*D* | **X**, *G*_*s*_, *A*_*s*_ = *a*, *θ*_*a*_) is a logistic regression of the disease status indicator, *D*, on the covariates, **X**, and SNP genotype, *G*_*s*_, and with coefficient vector *θ*_1_ under H_a_ and is a logistic regression *D* on **X** with coefficient vector *θ*_0_ under H_o_. We place independent normal mean 0, standard deviation 10 prior distributions on all components of *θ*_0_ and *θ*_1_, with exception of the coefficient of *G*_*s*_, which is accorded a normal mean 0, standard deviation 0.25 prior distribution, as the majority of log–odds estimates cited in the GWAS catalog are smaller than 0.5 in absolute value. We estimate the SNP–specific marginal likelihoods under each hypothesis of association using the Laplace approximation (Kass and Raftery, 1995) implemented in software described in Wilson et al. (2010) and available from the authors.

Since it is not always convenient or possible to directly calculate Bayes factors, we consider the performance of our method when applied to Bayes factors estimated from p-values using the approximation described in Sellke et al. (2001). These authors show that the Bayes factor *against* association can be approximated by the function −*ep*_*s*_ log_*e*_(*p*_*s*_) when *p*_*s*_ < 1/*e* and 0.0 otherwise, where *p*_*s*_ is a p-value from a standard test of association, and that this function provides a lower bound for that quantity that is sharp (i.e. accurate) for *p*_*s*_ < 1/*e*. As a consequence, its multiplicative inverse provides a sharp upper bound on the Bayes factor *in favor of* association. We evaluate our method using both this and the ratio of Laplace approximations described above to calculate **BF**_*s*_ in Equation 1.

### 4.2 Construction of Functional Signatures

In what follows, we detail the steps we took to assemble the case–control study of SNPs used to build and evaluate the models for a variant’s association status given the functional data. These comprised identification of a representative set of phenotype–associated SNPs to serve as ‘cases’ in the analysis and a matched set of ‘controls’ and the collection of a set of measurements related to function to serve as annotations for the variants. The process is depicted in Figure 4 and described in detail below.

**Figure 4.**
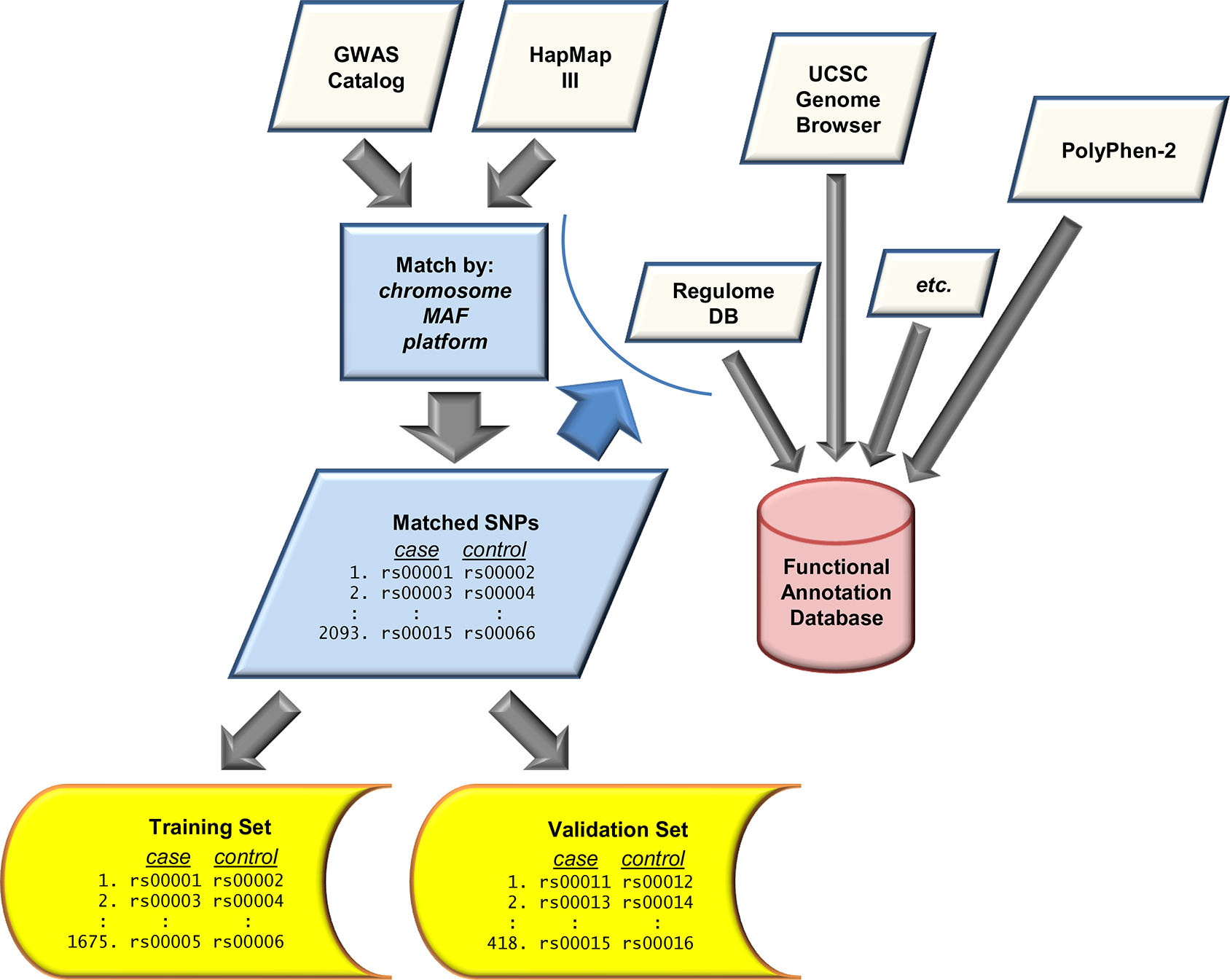
Construction of data sets and functional annotation database. Case SNPs from the GWAS Catalog are matched with control SNPs from HapMap III to generate training and validation sets. The matched SNP IDs and their locations are used to interrogate several online databases. These results are merged to build the functional annotation database.

#### Sampling Frame

Many genomewide genotyping arrays were designed to over–sample variants with characteristics related to their ability to explain phenotypic variation, such as proximity to coding regions, type of variant (e.g. missense) and minor allele frequency (MAF). Hence, a comparison of case SNPs identified using such assays to control SNPs drawn randomly from the HapMap or dbSNP, for example, may lead to spurious associations between assay design variables and the SNP’s association with a human phenotype. In order to avoid confounding due to the selection method employed in the design of the genomewide genotyping platform, we constructed a sampling frame of SNPs by combining SNPs on the Affymetrix GeneChip Human Mapping 500 K Array Set and the Illumina HumanHap550 Genotyping BeadChip, as this generation of arrays and their predecessors cover most of the reported findings in the GWAS Catalog attributed to an Affymetrix or Illumina product and these products were the most commonly used. We labeled SNPs in the sampling frame according to whether they appeared only on the Affymetrix list, only on the Illumina list or on both and confined attention to those variants appearing in both the Genome Browser’s dbSNP 130 and HapMap Release 27 tables (see Supplemental Table 1) and having a MAF estimated in HapMap’s CEU sample to be 0.05 or larger. The sampling frame comprises 803,991 SNPs with 421,072 unique to Illumina, 305,672 to Affymetrix and the remaining 77,247 common to both.

#### Case and Control Selection

The GWAS Catalog is subject to constant update and versions are available from several locations. We downloaded the GWAS Catalog from the Genome Browser (time stamp and location in Supplemental Table 1). We confined attention to non–CNV variants in the GWAS Catalog discovered by association studies utilizing an Affymetrix and/or an Illumina genomewide array and present in the sampling frame. We randomly chose a single representative of each set of SNPs appearing multiple times in the GWAS Catalog or sharing one or more ‘LD partners’ (see below). This left 2093 unique case SNPs, 1306 of which were unique to Illumina, 403 unique to Affymetrix and 384 in common. We randomly matched one control SNP drawn from the sampling frame to each case SNP on chromosome, platform (Illumina only, Affymetrix only, on both) and MAF rounded to the nearest 0.02. We excluded SNPs in the sampling frame in LD (R^2^ > 0) with one or more case SNPs as reported in the HapMap Release #27 LD files (see Supplemental Table 1) or sharing an LD partner with another control SNP.

#### LD Partner Identification

SNPs in the GWAS Catalog are arguably more likely to tag the variant that is directly associated with the phenotype than to be that variant (Hindorff et al., 2009). Hence, following Hindorff et al. (2009) we identified and annotated each case and control SNP’s ‘LD partners.’ We defined LD partners as those SNPs with R^2^ ≥ 0.8 with a case or control SNP as reported in the HapMap Release #27 LD files. Hindorff et al. (2009) chose a threshold of 0.9 but noted that their results were nearly the same when using thresholds of 1.0 and 0.8. We identified 20,924 LD partners of the case SNPs and 23,779 LD partners of the control SNPs.

#### Annotation Data

All data drawn from the UCSC Genome Browser (Rhead et al. (2010)) used the “March 2006 (NCBI36/hg18)” assembly. Supplemental Table 1 provides locations, revision dates and references for each of the annotation files referred to below. In what follows, we describe each class of annotation variable, its source and the parameterization we use for it in the models we fit.

Variants described in dbSNP (Sherry et al., 2001) release 130 are classified according to their predicted function as determined by their locations relative to known genes in the reference assembly. Variants that fall within the coding sequence of a known gene are further described as ‘non–synonymous’ if they result in a change to the associated amino acid or ‘synonymous’ if they do not. A variant may have several such designations; for purposes of our analysis, we confine attention to each variant’s primary designation. Those observed among the SNPs included in our analysis are ‘unknown,’ ‘coding–synon,’ ‘intron,’ ‘near–gene–3,’ ‘near–gene–5,’ ‘nonsense,’ ‘missense,’ ‘untranslated–3,’ and ‘untranslated–5.’ Given the small number (*n* = 5) of nonsense variants, we created a ‘coding–nonsynonymous’ designation by combining the ‘missense’ and ‘nonsense’ categories; similarly, we combined the ‘untranslated–3,’ and ‘untranslated–5’ designations into the category ‘untranslated.’

Measures of sequence conservation are frequently employed as evidence regarding the disease association status of rare missense variants (Tavtigian et al., 2008). We examined the PhyloP evolutionary conservation scores of Siepel et al. (2006) applied to 28– and 44–species alignments, and to those alignments restricted to the placental mammals and human, for their ability to predict the disease association status of common variants. Each of the four relevant Genome Browser tables provides the sum of the score, its sum of squares and the number of nucleotides that contribute to these statistics within ranges of contiguous nucleotides. We calculated a standardized score (mean divided by standard deviation) for each range and alignment and assigned these values to SNPs within the range. The PhyloP conservation scores exhibited pairwise correlations of up to 0.984. The top four, six, and nine PCs explain 90%, 96%, and 99% of the variability, respectively, in the 24 variables. The top four PCs were used.

The Database of Genomic Variants (DGV; Iafrate et al. (2004); Zhang et al. (2006)) is a compilation of reported genomic alterations spanning more than 1000 bases (>100 in the case of indels) observed in healthy subjects. We formed three variables indicating, respectively, whether (=1) or not (=0) each SNP falls in a region of copy number variation, a region containing insertions or deletions or a region with inversions.

The ENCODE Project (The ENCODE Project Consortium, 2007, 2011) is an ambitious project to identify and characterize the various functional elements present in the human genome sequence and to facilitate public access to the data it generates; its overarching objective is to improve our knowledge of human disease processes by providing a more comprehensive understanding of human molecular biology. Application of ENCODE functional annotation data to the design, analysis and interpretation of GWAS studies is one way in which ENCODE data can quickly be put to use to shed light on human disease processes (The ENCODE Project Consortium, 2011). To this end, we examine the utility of the recently released ENCODE regulation supertrack data available from, and displayed on, the Genome Browser for *a priori* prediction of functional, disease-associated variants. In particular, we include variables (see below) summarizing: transcription levels assayed in six cell lines by RNA–seq (Mortazavi et al., 2008; Langmead et al., 2009) and represented as a density measure of signal enrichment (’raw signal’); density measures of signal enrichment for H3K4Me1 (Histone H3 Lysine 4 monomethylation) associated with enhancer and promoter activity measured in eight cell lines, similarly coded measures of promoter– and enhancer–associated H3K27Ac (Histone H3 Lysine 27 acetylation) in eight cell lines and of promoter–associated H3K4Me3 (Histone H3 Lysine 4 tri–methylation) in nine cell lines (Bernstein et al., 2006; Mikkelsen et al., 2007); evidence for the variant falling within a DNaseI hypersensitivity cluster (Sabo et al., 2006, 2004); and the evidence for transcription factor binding measured via ChIP–seq (Euskirchen et al., 2004, 2007; Martone et al., 2003; Robertson et al., 2007; Rozowsky et al., 2009).

The Broad ChIP–seq, Caltech RNA–seq, and PhyloP signal tracks are summarized at the level of genomic bins. The ChIP–seq signals are measured within 118,084 contiguous bins of 25,600 bases apiece. The RNA–seq and PhyloP signals are measured in sets of non-overlapping, non-uniform bins. Bins are indexed according to the hierarchical scheme described in (Kent et al., 2002).

The ENCODE database provides basic summary statistics (the minimum, range, count, sum and sum of squares) of the signal enrichment density measures within each bin. For purposes of our analysis, we summarized each cell line’s bin level data by log_e_(sum/count), log_e_(maximum), and log_e_(z), where z is the standardized score; in bins where sum = 0, we set sum = 1 (sums are non–negative and range across six orders of magnitude).

The three Broad types (signal enrichment for H3K4Me1, H3K27Ac and H3k4Me3 histone modifications) comprise data on eight, eight and nine cell lines, respectively. We found significant pairwise correlations among the 75 variables (25 each of log(mean), log(maximum), and log(z)), ranging as high as 0.949, and therefore conducted a principal components analysis to identify the linear combinations, i.e. principal components (PCs), that explain most of the variability in the data. The top 18, 27 and 44 PCs explain 90%, 95% and 99% of variability, respectively, in the 75 measures. Finally, we mapped each SNP to the appropriate Broad bin and annotated each with the top 18 PCs for purposes of the analysis.

The Caltech tables comprise RNA–seq raw signal enrichment data on six cell lines. In addition to the three ENCODE variables described above, an indicator variable for a SNP falling within a bin was included. Pairwise correlations among the 24 variables ranged as high as 0.970. The top 11, 12, and 16 PCs explain 92%, 95%, and 99% of the variability. The top 11 PCs were used in the analysis.

The transcription factor ChIP–seq data are summarized by scores, ranging from 6 to 1000, measuring strength of evidence for binding within specified, sometimes overlapping, chromosomal bins (‘clusters’). We summarize these data as they apply to each SNP using two variables: the number of clusters it intersects with (‘TFBSfreq’) and the average log_e_(score) (‘logTFBS’) assigned to those clusters (coded as 0 if the SNP does not intersect with a cluster). Similarly, the DNaseI hypersensitivity data are summarized by scores, ranging from 16 to 1000, within specified chromosomal bins (‘clusters’). We summarize these data as they apply to each SNP using (1) an indicator for the variant falling within a clusters and (2) the log_e_(score) assigned to that cluster.

The Open REGulatory ANNOtation database (ORegAnno) Montgomery et al. (2006); Griffith et al. (2008) is a curated collection of regulatory elements. The Genome Browser ORegAnno table provides start and stop coordinates and annotations for elements in the database. For purposes of our analysis, we summarize these data with an indicator variable for whether or not a variant falls within an ORegAnno regulatory region.

PolyPhen–2 (PPh2; Adzhubei et al. (2010)) assigns to nonsynonymous SNPs a probability of being damaging based on the sequence, phylogenetic and structural information characterizing the amino acid substitution.

RegulomeDB (Boyle et al., 2012) annotates SNPs with known and predicted regulatory elements in the intergenic regions of the human genome. Each SNP is assigned one of seven categories based on its likelihood of affecting protein binding.

#### Model

For purposes of the analysis, we assumed that blocks and the SNPs within the blocks were independent conditional on the functional data. We modeled the probability that a SNP *s* in block *b* was an associated SNP, *π*_*sb*_, given the functional data for that SNP, *F*_*sb*_, using the logisticregression model logit(*π*_*sb*_) = *α* + F_sb_*β*. We assumed that there was at least one associated SNP in each case block and that there were no associated SNPs in control blocks. Hence, each case block contributed the factor 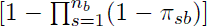 to the likelihood, while each control block contributed 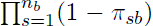. As a result, we expected *at least* 1,675 of the 48,888 SNPs in the training set to be phenotype associated. This corresponds to *α* = −3.34 (columns of *F* are centered); if 10% (alt 20%) of case blocks contain two phenotype associated SNPs, *α* = −3.24 (alt −3.15). Hence the normal mean −3.24, standard deviation 0.1 prior distribution we placed on *α* is consistent with our expectation that there were fewer than 2,178 (= 1.3 × 1,675) true phenotype associated variants among the case blocks.

Our specification of the prior distribution on *β* was guided by the observation that, in the normal model with the normal–exponential–gamma (NEG) distribution as prior on the mean and the variance known, the posterior mode is identically zero when the maximum likelihood estimator (MLE) is in a neighborhood around zero, but rapidly converges to the MLE as the MLE diverges from zero (this setting approximates the more general one in which the NEG distribution is used as the prior distribution for a parameter whose likelihood is approximately normal). The NEG distribution is specified by its shape and scale parameters and the width of the threshold neighborhood is a function of these parameters. For purposes of our analysis, we chose parameter values for which no more than 10% of the coefficients are outside of the threshold region with probability 0.90, *a priori*. We placed independent NEG prior distributions on the components of *β*; in addition, we also considered the model with independent standard normal distributions on the components of *β*. Inference for each of these models was carried out using the training set and were evaluated using the evaluation set.

We used random–walk Markov Chain Monte Carlo (MCMC) algorithms (Metropolis et al., 1953; Gilks et al., 1996) to estimate summaries of the posterior distribution under each of the models. We started 10 independent chains per model from starting points drawn from the prior distribution. In each case, step sizes were adusted so that parameter level acceptance ratios fell between 0.3 and 0.5 during an initial, ‘burn–in’ set of iterations not used for inference. We fixed the step sizes and ran the 10 chains from their leave–off positions for an additional 50,000 iterations per chain. Inspection of trace plots, as well as computation of the Gelman–Rubin (Gelman and Rubin, 1992), Heidelberger–Welch (Heidelberger and Welch, 1983), Raftery–Lewis (Raftery and Lewis, 1996), and Geweke (Geweke, 1992) diagnostics implemented in the CODA package (Plummer et al., 2010) in R (Ihaka and Gentleman, 1996), indicated satisfactory convergence. We thinned the 10 chains by 1,000 and combined them to produce a sample of 500 coefficient vectors.

We used the concordance index (CI) to measure the out–of–sample predictive accuracy of the model. We calculated the CI as the fraction of matched pairs in the ‘evaluation set’ in which the average probability of association given the functional data over the *n*_1_ SNPs in the case block (‘*b*1’) was larger than the corresponding average over the *n*_0_ SNPs in the matched control block (‘*b*0’); i.e. if

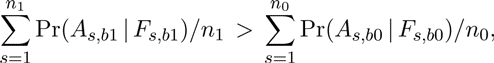

where we estimated Pr(*A*_*s*,*bn*_ | *F*_*s*,*bn*_) by

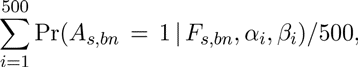

where the *α*_*i*_ and *β*_*i*_ are MCMC samples saved from analysis of the training data.

### 4.3 Evaluation

We carried out a genome–wide association analysis of serous ovarian cancer using the methods described above. The data for this analysis were drawn from GWAS studies conducted in the US (Permuth-Wey et al., 2011) and the UK (Song et al., 2009). The genotype data from these studies were combined and imputed to HapMap III density, resulting in an data set comprising analyzable genotypes at 2,500,004 SNPs for 7,272 subjects of European ancestry. The association analysis was confined to the 2,004 cases with advanced stage serous ovarian cancer and the 3,272 available controls and was adjusted for study site and the first two principal components of the sample genotypes. We calculated Bayes factors (BFs) as described above and set the prior probability of association to be 0.00001 when estimating posterior probabilities; ranks are invariant to this choice. P-values used in the Bayes factor approximation were from likelihood ratio tests of the model including versus the model excluding a SNP.

Several large scale studies conducted to follow up promising associations from these GWAS have identified the eleven genome–wide significant loci listed in Table 4. We treat these as established or ‘true positive’ associations for purposes of evaluating the various association measures. In addition, we identified a set of likely unassociated, ‘true negative’ SNPs from among 22,254 GWAS followup SNPs placed on the iCOGS chip (Pharoah et al., 2013). This analysis included 8,344 cases with advanced stage serous ovarian cancer and 22,913 controls of European ancestry and was adjusted for study site and the first five European ancestry principal components. We identified a subset of 5,155 SNPs with strong evidence *against association* (defined as BF < 0.1 on Jeffreys’ scale of evidence (Jeffreys, 1961)) to serve as the ‘true negatives.’

We compared the rankings of these two sets of SNPs in the original GWAS analysis when association was measured using genotype data only to those obtained with incorporation of the functional signatures. We compared the procedures based on their power to identify the truly associated variants for follow–up assuming budgets allowing for evaluation of the top 5,000 or 10,000 SNPs.

In most association studies, genotypes are determined, through a combination of genotyping and imputation, for only a subset of the universe of variants. In this setting, it is standard to rely on correlations between genotyped variants (‘tags’) and those that are ‘tagged’ (not genotyped) to identify and localize associations. Likewise, the utility of functional signatures in a typical study will depend on the degree to which they reflect the likelihood of function of both the tag for which it is calculated and for the set of variants it tags. We evaluated correlations between functional signatures, defined as (*F*_*s*_*β*), for adjacent pairs of SNPs included in the ovarian cancer GWAS analysis. We identified the quantile of each adjacent variant pair in the overall distribution of distances measured in base pairs (BPs). For purposes of this analysis, we defined quantiles in increments of 0.025, i.e. with each containing 2.5% of the mass of the distance distribution. We estimated the Pearson correlation between the functional signatures of the adjacent SNP pairs within each quantile and plotted these estimates against BP distance, locating the estimates at the midpoints of the quantile bins.

## Data Access

The data used to construct and evaluate the functional signatures we describe are available at ftp://stat.duke.edu/pub/Users/iversen/FunctionalSignatures/.

## Acknowledgements

**Funding:** This work was funded by the National Institutes of Health through its Genes and Environment Initiative, grant number R01–HL090559 and through R21–ES020796 (NIEHS) and by the National Science Foundation via DMS–1106891. The ovarian cancer multi–GWAS analyzed herein were provided by the FOCI project (1U19-CA148112) of NCI’s GAME–ON consortium.

## Supplement

**Supplemental Table 1.**
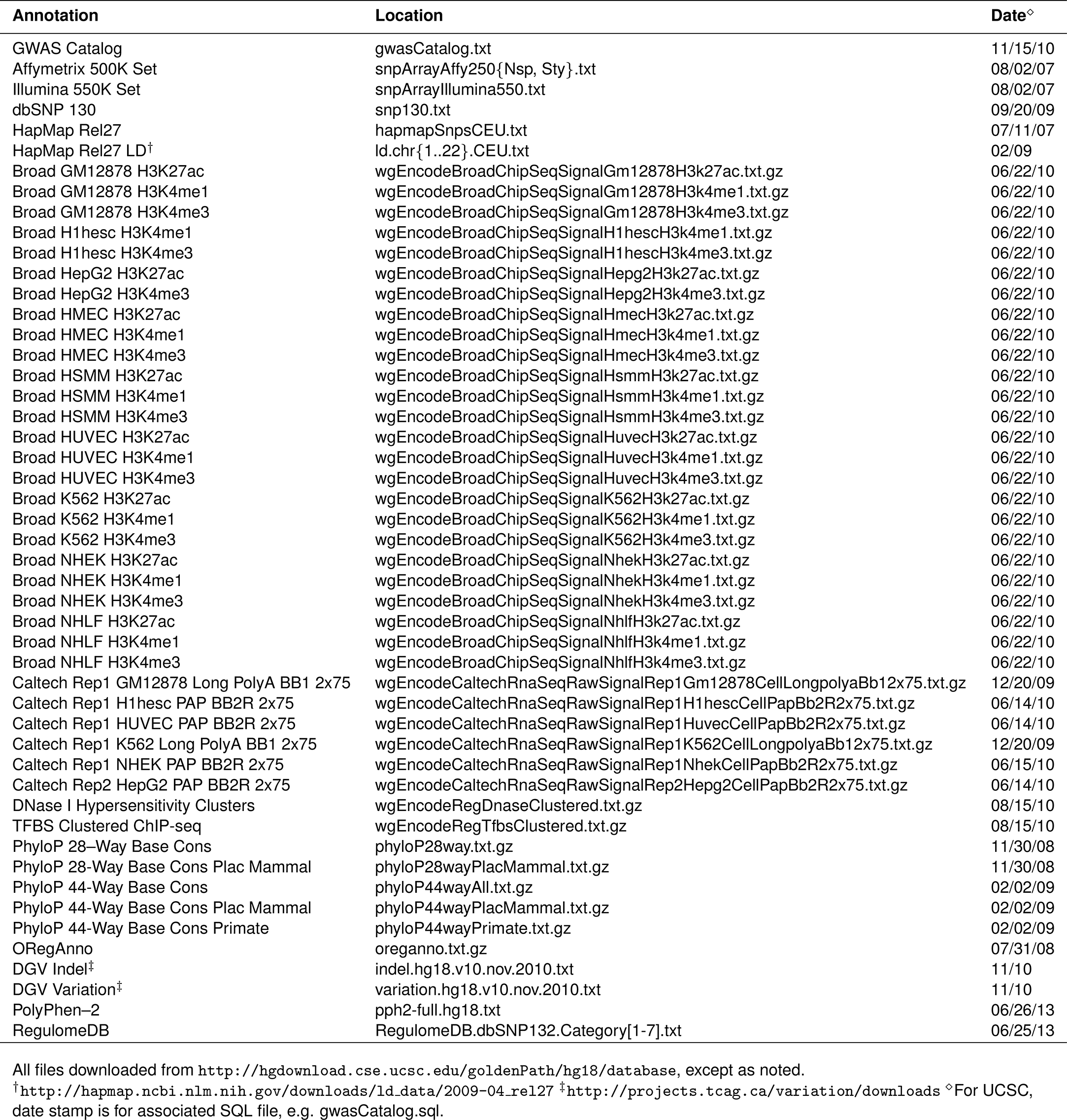
Summary of data sources.

**Supplemental Figure 1.**
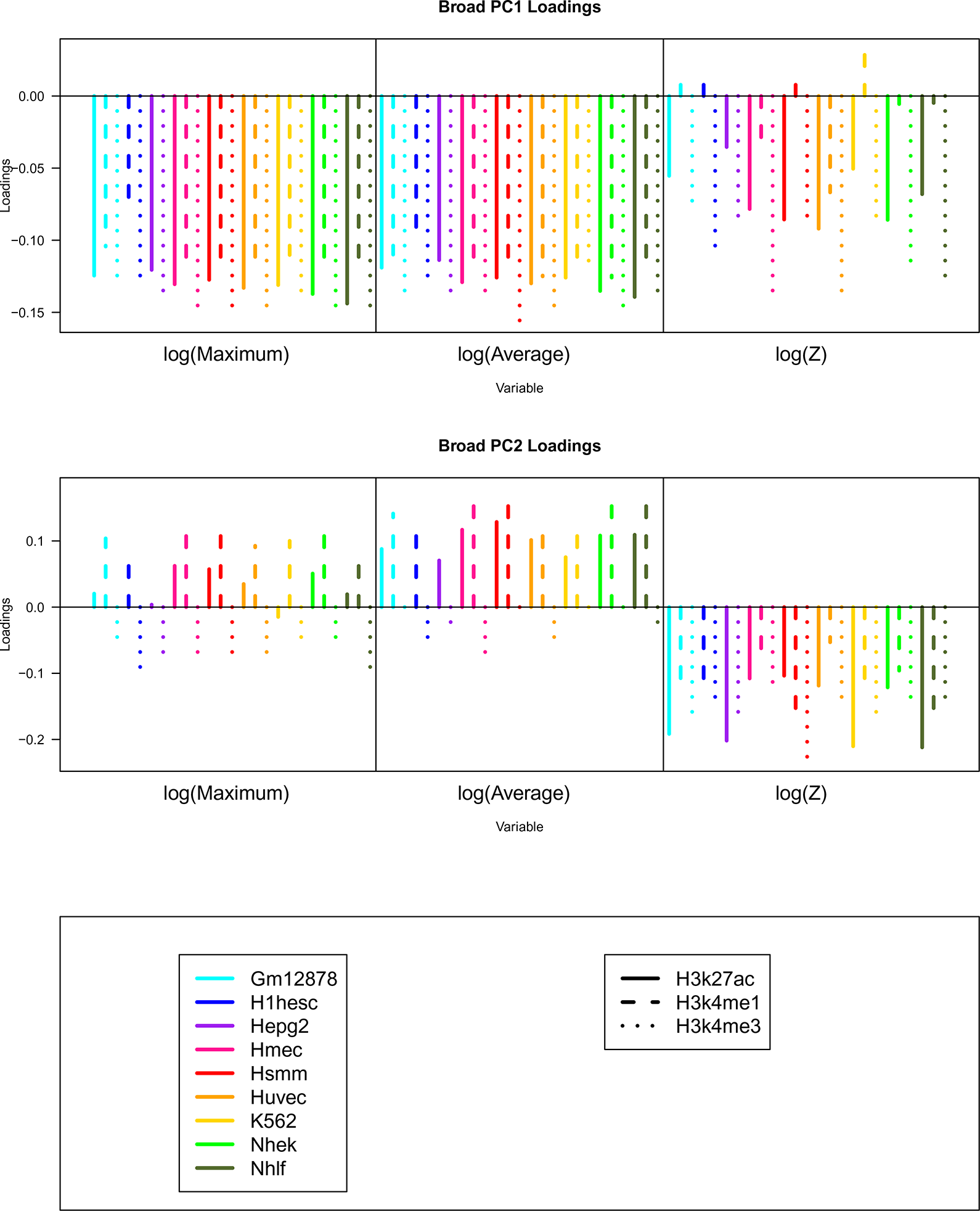
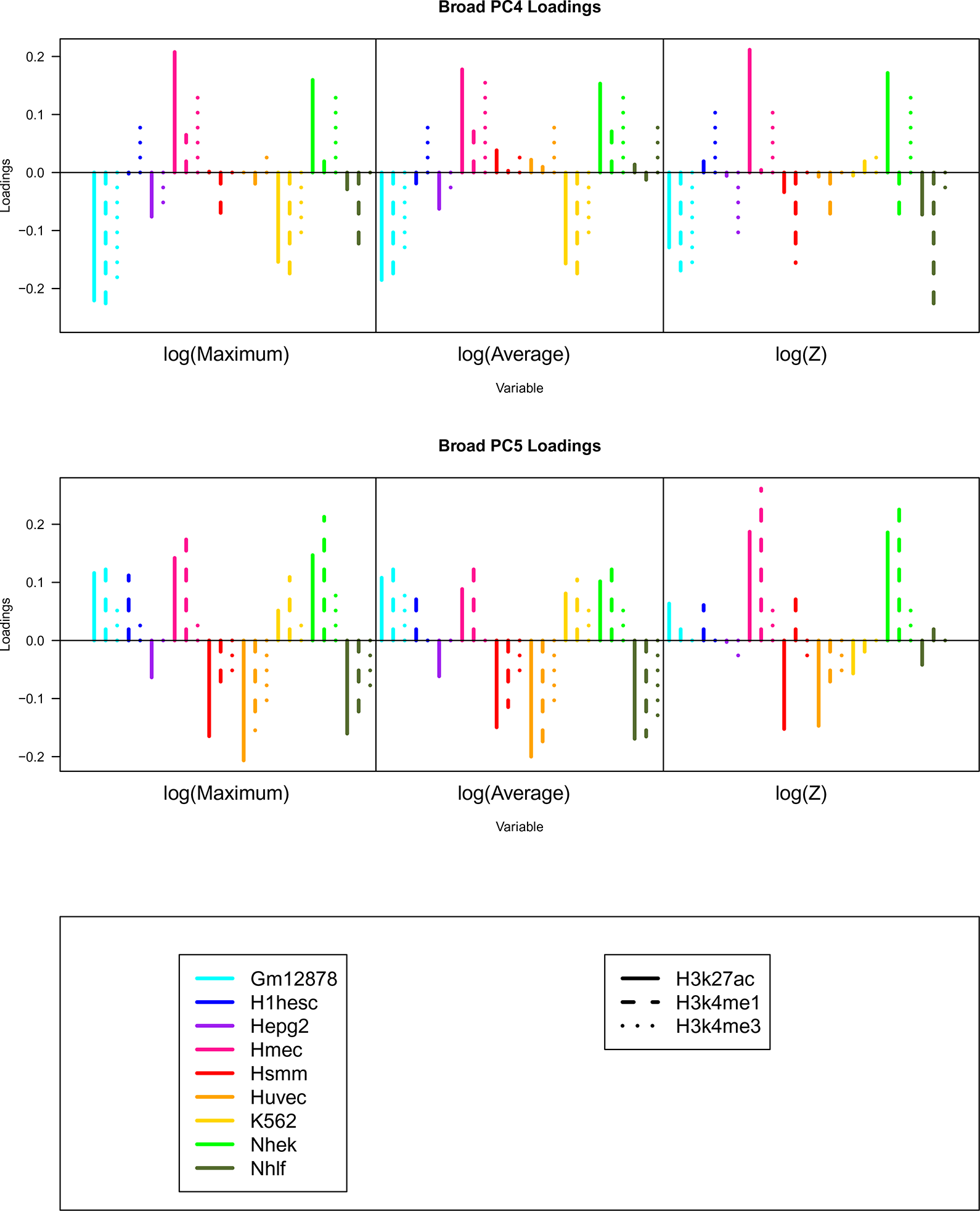
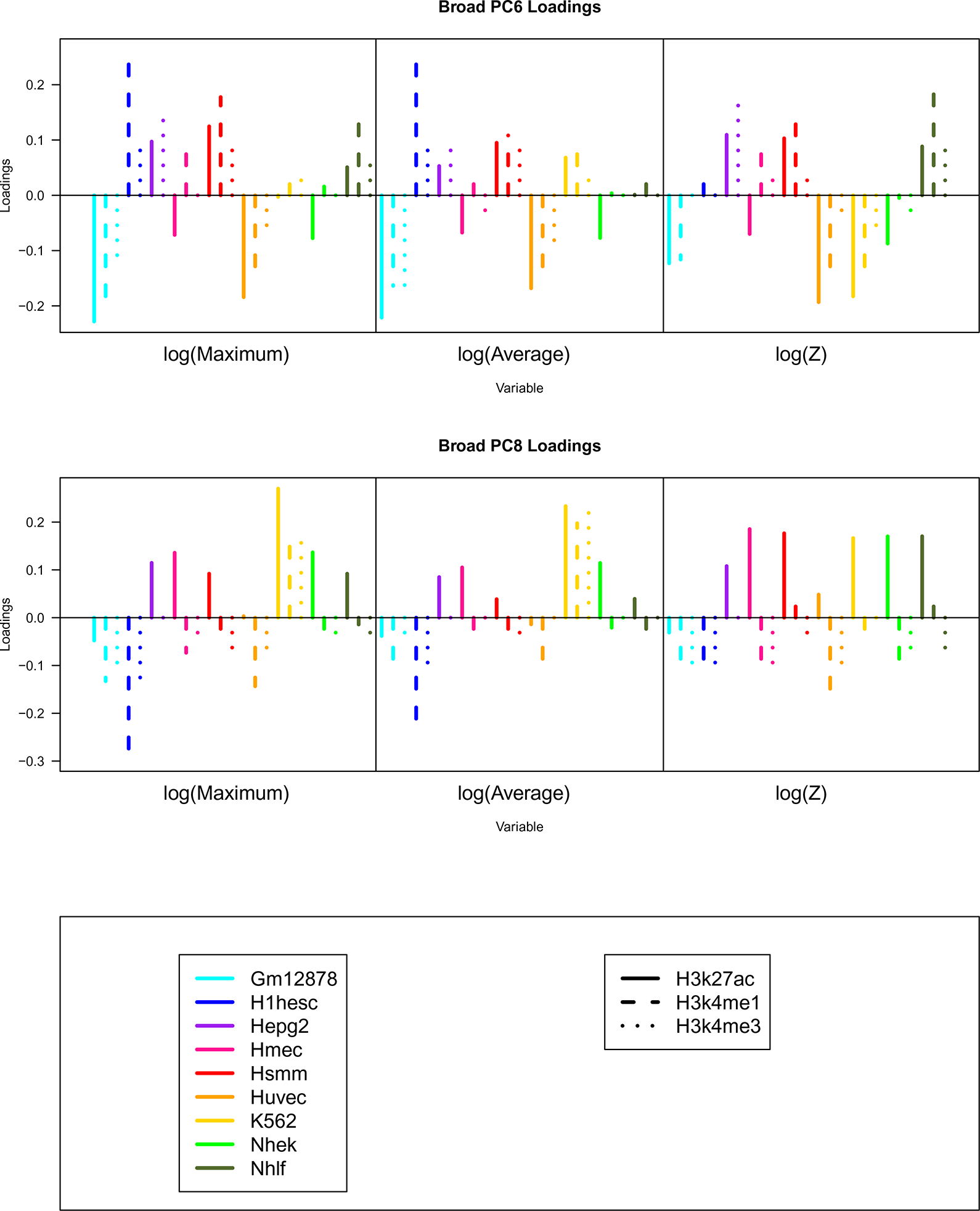
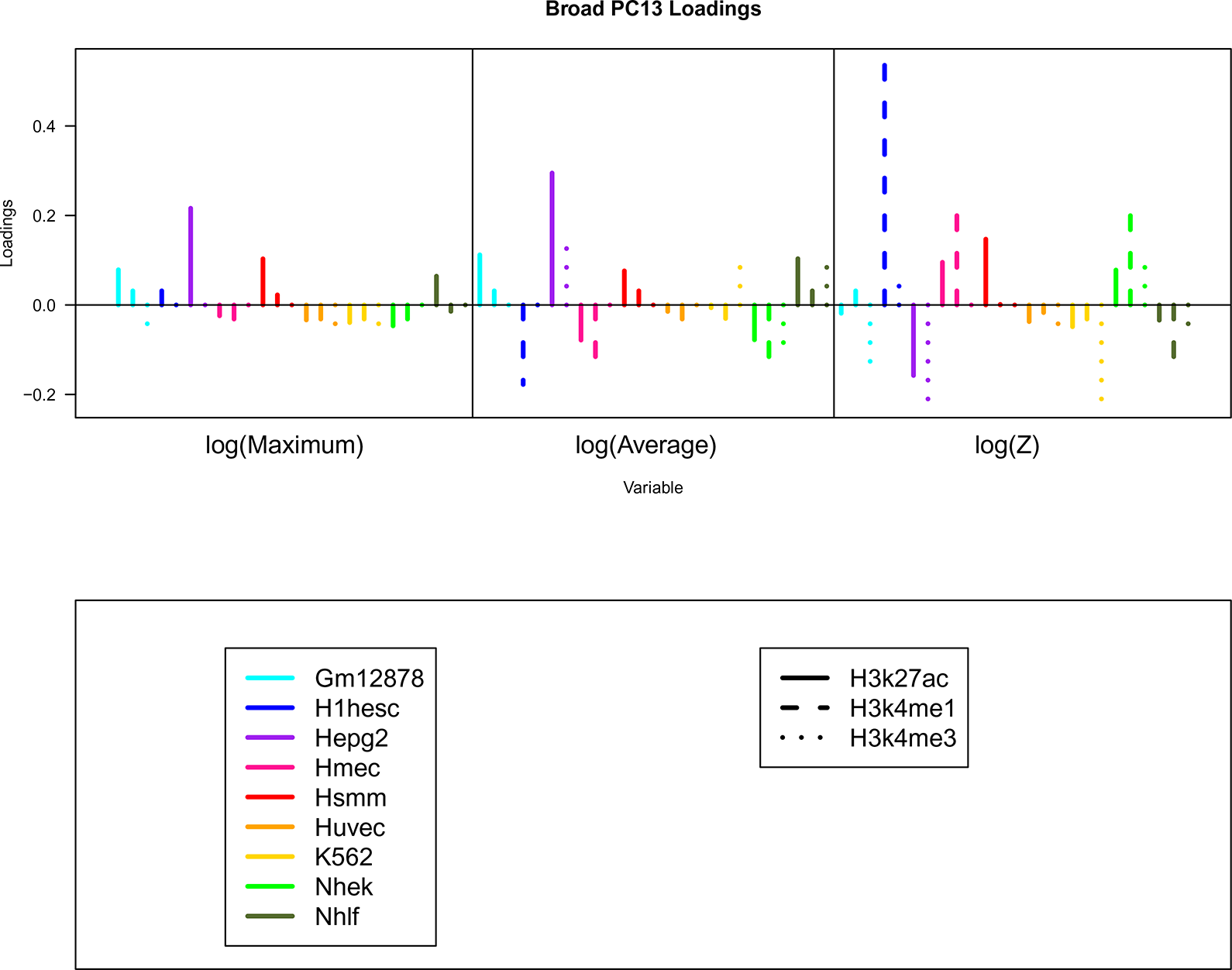
Plot of the loadings (weights in the linear combinations) for the most highly associated Broad promoter/enhancer ChIP–seq principal components (PCs) as they depend on histone modification, cell line and summary statistic.

**Supplemental Figure 2.**
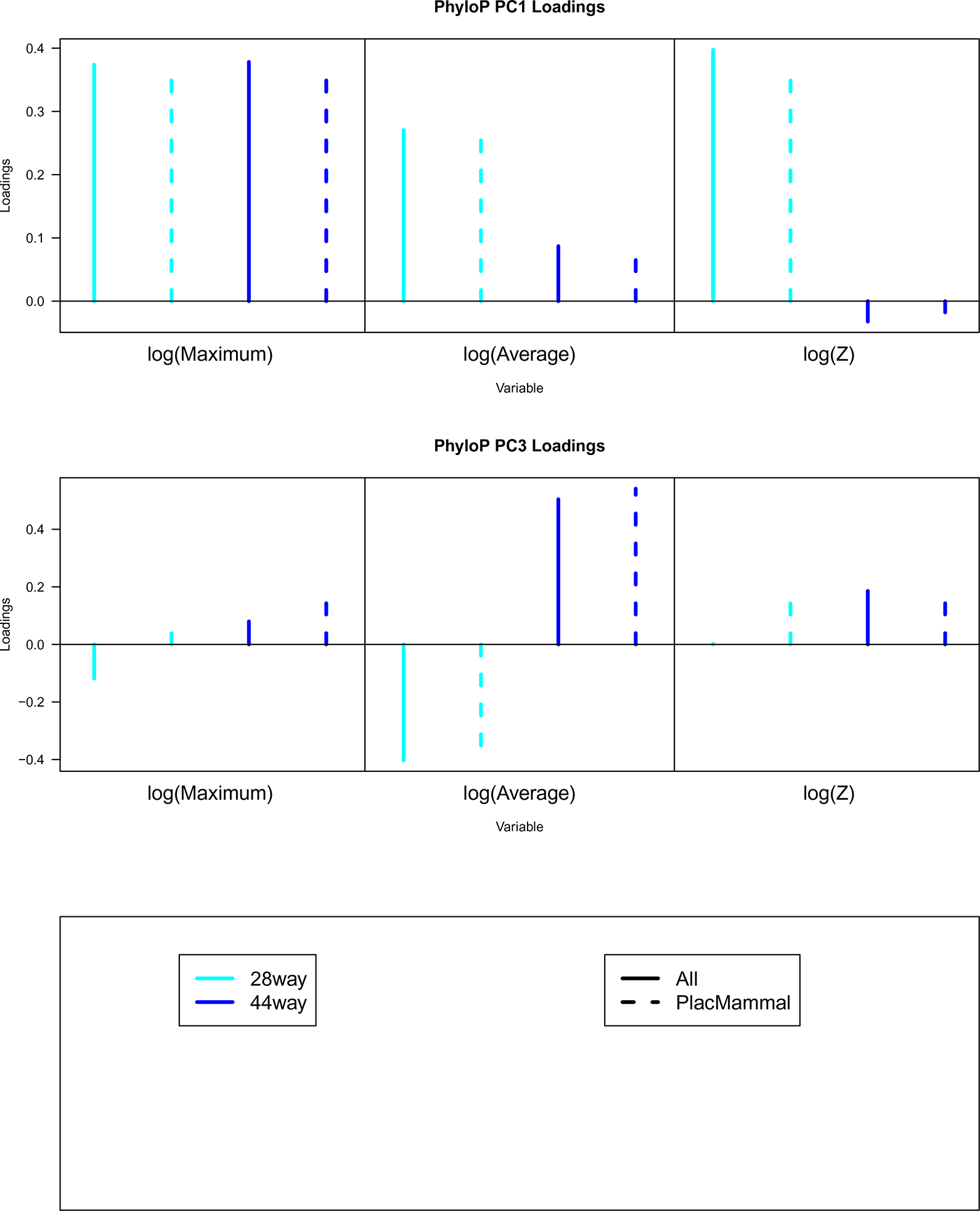
Plot of the loadings (weights in the linear combinations) for the most highly associated sequence conservation principal components (PCs) as they depend on number of species, depth of alignment and summary statistica.

**Supplemental Figure 3.**
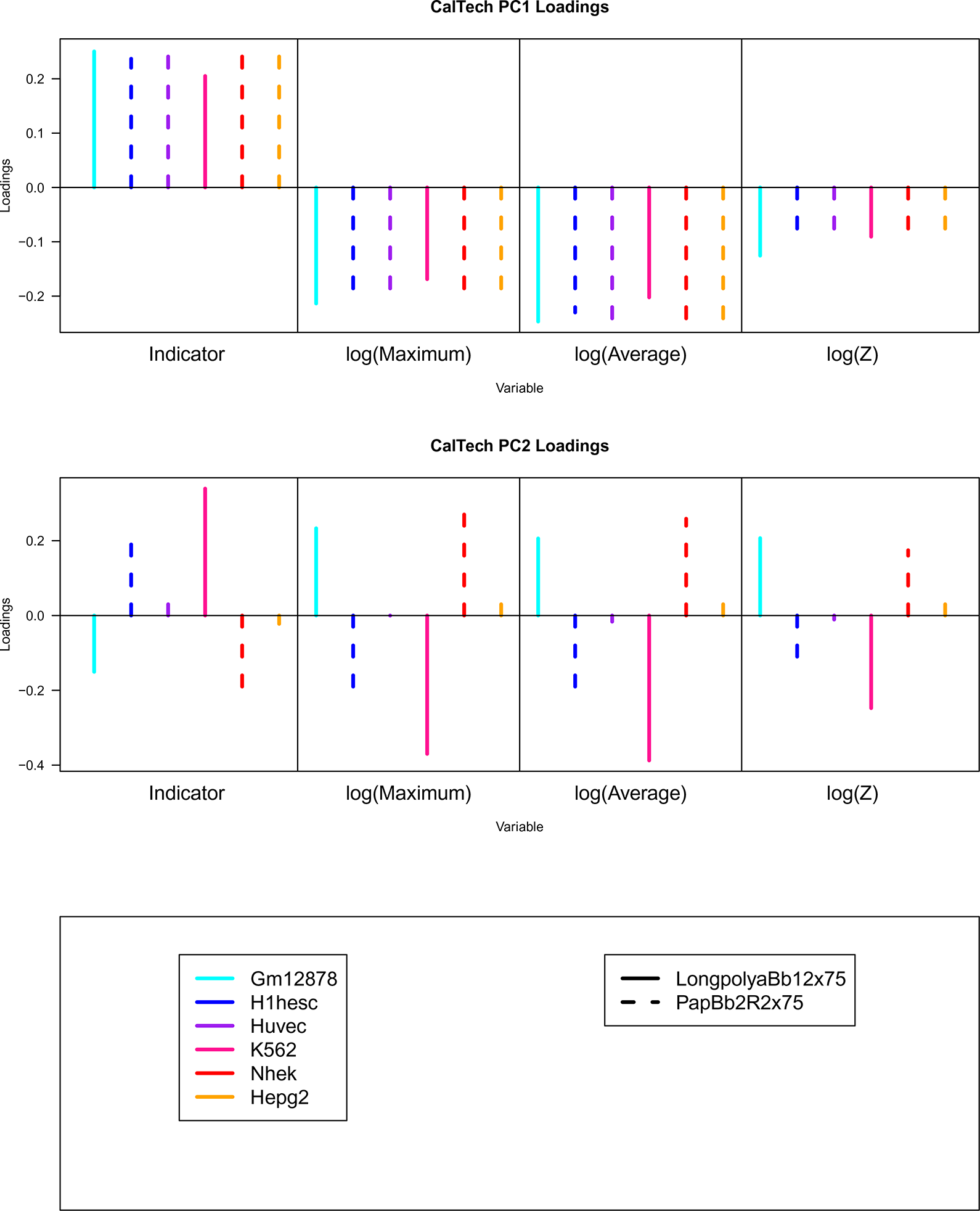
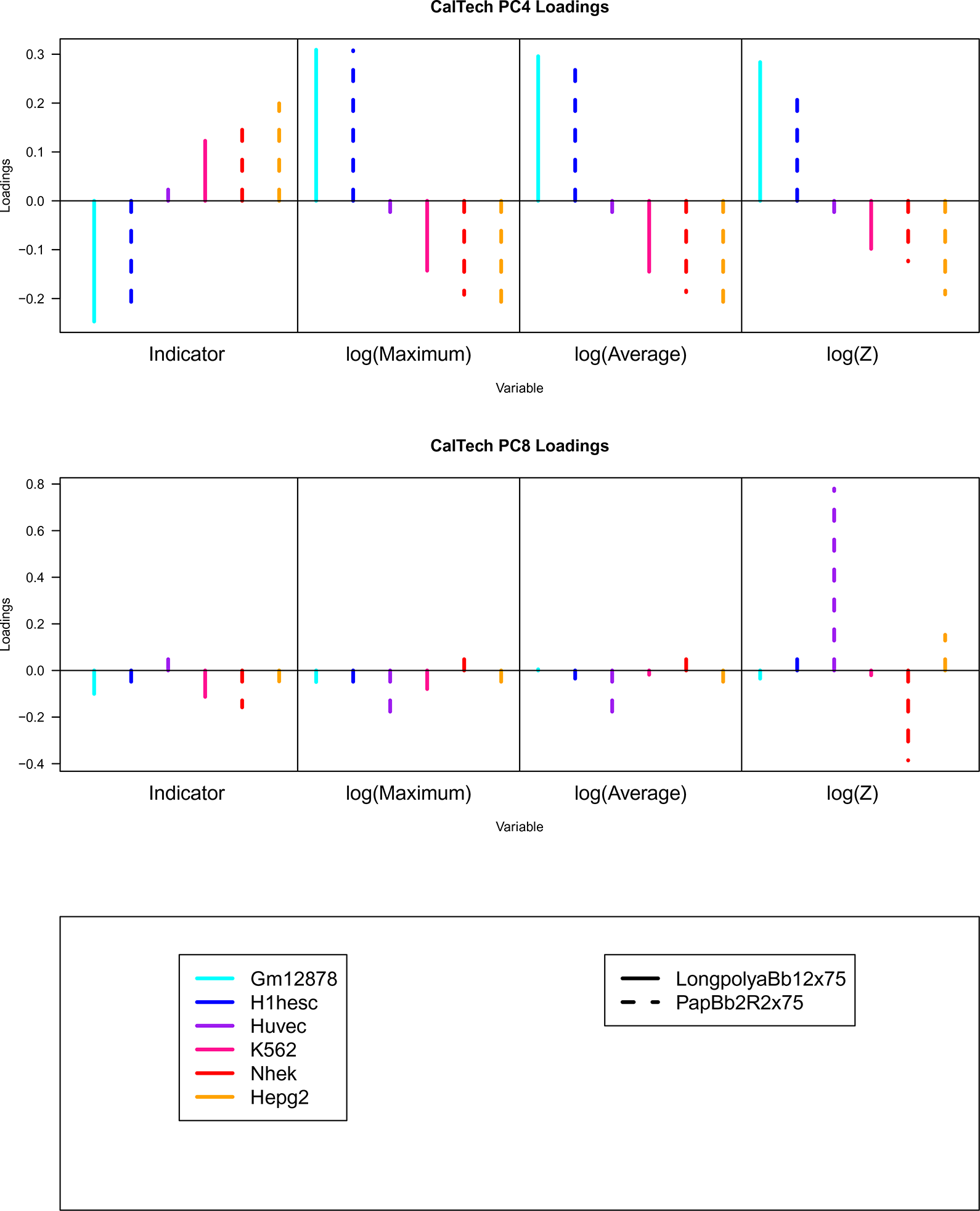
Plot of the loadings (weights in the linear combinations) for the most highly associated CalTech RNA–seq principal components (PCs) as they depend on cell line and summary statistic.

